# *Trypanosoma brucei* Pex13.2 is an accessory peroxin that functions in the import of PTS2 proteins and localizes to subdomains of the glycosome

**DOI:** 10.1101/474080

**Authors:** Logan P. Crowe, Kathleen R. Nicholson, Christina L. Wilkinson, Meredith T. Morris

## Abstract

Kinetoplastid parasites including *Trypanosoma brucei, Trypanosoma cruzi* and *Leishmania* harbor unique organelles known as glycosomes, which are evolutionarily related to peroxisomes. Glycosome/peroxisome biogenesis is mediated by proteins called peroxins that facilitate organelle formation, proliferation and degradation, and import of proteins housed therein. Import of matrix proteins occurs via one of two pathways that are dictated by their peroxisome targeting sequence (PTS). In PTS1 import, a C-terminal tripeptide sequence, most commonly SKL, is recognized by the soluble receptor Pex5. In PTS2 import, a less conserved N-terminal sequence is recognized by Pex7. The soluble receptors deliver their cargo to the import channel consisting minimally of Pex13 and Pex14. While much of the import process is conserved, kinetoplastids are the only organisms to have two Pex13s, *Tb*Pex13.1 and *Tb*Pex13.2. In previous studies, GFP-tagged *Tb*Pex13.1 localized to glycosomes and silencing either protein in the stage of the parasite that lives in the mammalian bloodstream impaired glycosome protein import and slowed parasite growth. While these findings suggest Pex13s are involved in protein import, the mechanisms by which they function are unknown and it is unclear why kinetoplastids would require two Pex13s. In this work, we demonstrate that *Tb*Pex13.2 is associated with the glycosome membrane with its N-terminus facing the cytoplasm. Super-resolution microscopy reveals that *Tb*Pex13.2 localizes to a few (1-3) foci per glycosome and import of PTS2 proteins was disrupted in *Tb*Pex13.2-deficient cells suggesting it may be an accessory factor for PTS2 import.

## INTRODUCTION

The family Kinetoplastea includes a number of medically relevant parasites including *Trypanosoma brucei, Trypanosoma cruzi* and several species of *Leishmania* that cause human African trypanosomiasis, Chagas disease, and several forms of leishmaniasis. Treatments for these diseases are expensive, toxic, and difficult to administer. Additionally, the growing resistance to current therapies necessitates the need for the identification of new drug targets.

All kinetoplastids have unique membrane-bounded microbodies called glycosomes that are evolutionarily related to peroxisomes of higher eukaryotes (*1*, *2*). Peroxisomes and glycosomes lack DNA and import most matrix proteins post-translationally (*2*). Glycosomes are unique in that they harbor many of the enzymes in the glycolytic pathway, which are cytosolic in higher eukaryotes (*3*–*5*). The essential nature of these organelles makes them a good drug target. Highlighting the druggable nature of these organelles, recent work has shown that small molecules, which interrupt protein import into glycosomes, were lethal to *T. brucei* (*6*).

Peroxisome biogenesis is regulated by proteins called peroxins (PEXs) that govern organelle formation, proliferation and degradation, as well as protein import (*2*, *3*, *7*). Import of proteins into peroxisomes involves binding of soluble receptor proteins, either Pex5 or Pex7, to a targeting sequence in the cargo protein (*8*, *9*). Pex5 binds to a C-terminal tripeptide with the consensus sequence of SKL called a peroxisome targeting sequence 1 (PTS1) while Pex7 binds to a less conserved N-terminal sequence termed PTS2 (*2*, *10*, *11*). The receptor-cargo complex then docks at the peroxisome membrane through interactions with the glycosome membrane proteins Pex13 and Pex14, which make up the import channel (*12*). After import, the receptors are recycled via a ubiquitination process involving Pex2, 10, and 12 (*13*).

While much of the import pathway is conserved, kinetoplastids are unique in that they have two Pex13s, which have been designated *Tb*Pex13.1 and *Tb*Pex13.2 (*14*, *15*). These proteins share low sequence identity with each other or with Pex13s from higher eukaryotes. In previous studies, *Tb*Pex13.1, which was the first identified, localized to glycosomes. Silencing of the protein in the mammalian bloodstream (BSF) and insect stage, procyclic form (PCF) parasites yielded parasites with glycosome protein import defects and slowed growth rates (*15*). Later, iterative database searches resulted in the identification of *Tb*Pex13.2 (*14*). Silencing of this second Pex13 via RNA interference in BSF parasites resulted in mislocalization of *Tb*Pex14 and *Tb*aldolase and a defect in growth rate. Prior to our work, *Tb*Pex13.2 RNAi cells lines could not be established in PCF parasites (*14*).

In pursuit of understanding function, we confirmed that native *Tb*Pex13.2 is an integral glycosome membrane protein and demonstrate that its N-terminus is on the cytoplasmic side of the membrane. Super-resolution microscopy reveals that *Tb*Pex13.2 localization is restricted to a few foci (1-3)/organelle and *Tb*Pex13.2-deficient PCF parasites exhibit a defect in PTS2 import, suggesting that it may serve as a co-receptor in PTS2 import.

## MATERIALS AND METHODS

### Cell culture and transfection of *T. brucei*

Procyclic form (PCF) 2913 and bloodstream form (BSF) 9013 expressing T7 polymerase and tetracycline (tet) repressor (*16*) were maintained in SDM79 (or the minimal glucose medium SDM79θ containing 5 μM glucose) (*17*) and HMI-9, respectively. Expression vectors for epitope-tagged proteins were generated by cloning the open reading frame of *Tb*Pex13.2 into the pXS2 (PCF) vector or pXS6 (BSF) vectors possessing either a blasticidin resistance or puromycin resistance gene (*18*). For transfection, 20 μg plasmid DNA was linearized (pXS2, pXS6: MluI; pZJM: NotI) and electroporated in 4 mm cuvettes (BioRad GenePulser Xcell; exponential, 1.5kV, 25μF). Twenty-four h after electroporation, culture media was supplemented with appropriate drug for selection: 15 μg/ml G418; 50 μg/ml hygromycin; 2.5 μg/ml phleomycin; 1 μg/ml puromycin; 10 μg/ml blasticidin. RNA interference (RNAi) cell lines were generated by cloning nucleotides 41-441 of *Tb*Pex13.2 into the inducible pZJM vector possessing dual opposing T7 promoters and a phleomycin resistance marker (*19*). Usually, we grow RNAi cell lines in tet-free media to reduce leaky expression from RNAi plasmids. However, this was not required for these cell lines as we did not observe leaky expression.

### *Tb*Pex13.2 antibody production

Polyclonal guinea pig antisera was generated against truncated recombinant *Tb*Pex13.2 (Thermo Scientific). Amino acids 2-150 of *Tb*Pex13.2 fused to an N-terminal His_6_ tag was expressed using the pQE30 expression system (Qiagen) and purified using a Ni-NTA column under denaturing conditions using 8M urea as described (*20*).

### Growth curves

Cells possessing the RNA inducible pZJM:*Tb*Pex13.2 vector were seeded at 10^5^ cells/ml in SDM79 (PCF) or 5×10^4^ cells/ml HMI-9 (BSF) and induced with 1 μg/ml doxycycline. PCF cells were allowed to grow to a density of 5×10^6^ cells/ml prior to passing back to 1×10^5^. BSF cells were allowed to grow to a density of 1×10^6^ cells/ml prior to passing back to 5×10^4^ cells/ml. Culture density was monitored by flow cytometry at 24 h intervals using an Accuri C6 flow cytometer (BD Biosciences).

### Sucrose gradient fractionation and western blot

Large-scale sucrose gradient fractionation was carried out as described previously (*21*). Small-scale fractionations for *Tb*Pex13.2 silencing were separated as described previously (*21*) with the following modifications: the post-nuclear lysate was separated on a 13 ml 20%-40% linear Optiprep gradient at 170,000*g* for 19 h in a Beckman SW-40Ti rotor at 4°C (acceleration 9 and deceleration coast), 500 μl fractions were taken from the top of the gradient and the protein concentration was determined by BCA assay (ThermoFisher). Protein from each fraction (2.5 μg) was separated by SDS-PAGE and analyzed by western blotting with antibodies against the glycosomal proteins: *Tb*aldolase (1:20,000), *Tb*Pex13.1 (1:10,000), *Tb*Pex13.2 (1:10,000), *Tb*Pex11 (1:4,000) provided by Dr. Christine Clayton (Zentrum für Molekulare Biologie der Universität Heidelberg, Germany) (*22*), *Tb*PFK (1:10,000) and *Tb*FBP (1:10,000) provided by Dr. Paul Michels (University of Edinburgh, UK), and the ER protein *Tb*BiP (1:100,000) provided by Dr. Jay Bangs (University at Buffalo, Buffalo, NY) (*18*).

### Protease protection assays

Protease protection assays were carried out using a modified protocol previously described (*22*). Cells (10^6^) were harvested at 800*g* for 10 min, washed once in PBS (150 mM NaCl, 1 mM KH_2_PO_4_, 5.6 mM Na_2_HPO_4_, pH 7.4), once in STE buffer (250 mM sucrose, 25 mM Tris-HCl, pH7.4, 1 mM EDTA), and resuspended in 98 μl ice cold STEN buffer (STE buffer supplemented with 150 mM NaCl) and 0.1 mM phenylmethylsulfonyl fluoride (PMSF). Cells were permeabilized with 2 μl 1mg/ml digitonin (final concentration of 0.02 mg/ml), vortexed for 5 sec and incubated at room temperature for 4 min. Following permeabilization, cells were centrifuged at 20,000*g* for 2 min and resuspended in 85 μl STEN buffer. Pellets were treated with either 10 μl water or Triton X-100 (1% v/v final) and either 5 μl water or 2 mg/ml proteinase K. Reactions were incubated on ice for 30 min and stopped by addition of 10% w/v trichloroacetic acid (TCA). Precipitated proteins were centrifuged at 17,000*g* for 10 min and washed once with acetone before being resuspended in cracking buffer (CB; 10% glycerol, 2% SDS, 2% ß-mercaptoethanol, 100mM Tris, pH 6.8, 0.1% bromophenol blue) and boiled at 100°C. Proteins were then analyzed by SDS-PAGE and western blotting.

### Membrane association assays

Membrane association assays were carried out as previously described by (*22*). For extraction of membrane proteins, 10^7^ cells were centrifuged at 800*g* for 10 min and resuspended in 300 μl of ice-cold low-salt buffer for 15 min (5 mM Tris-HCl pH 7.8, 1 mM EDTA, 0.1 mM PMSF, 4 μg/ml leupeptin). Cells were then passed through a pipette tip 10x and centrifuged at 20,000*g* for 30 min at 4°C. The insoluble pellet was resuspended in 300 μl high-salt buffer (25 mM Tris-HCl pH7.8, 0.5 M KCl, 1 mM EDTA, 0.1 mM PMSF, 4 μg/ml leupeptin) and incubated on ice for 15 min. After incubation, samples were centrifuged again at 20,000*g* for 30 min at 4°C. The insoluble pellet was resuspended in 300 μl 0.1 M Na_2_CO_3_ and incubated for 30 min on ice. Samples were then centrifuged at 120,000*g* for 1h at 4°C with a 500 μl cushion of 0.1 M Na_2_CO_3_, 0.25 M sucrose in a Beckman TLA100.3 rotor. Supernatant protein was precipitated by 10% w/v TCA and washed once with acetone before being resuspended in CB. Samples were then separated by SDS-PAGE and analyzed by western blot.

### Live-cell microscopy

Cells expressing either *Tb*AldoPTS2eYFP or *Tb*HKPTS2eYFP (Fig. 6B) were washed once with PBS, mounted on a slide and visualized using a Zeiss Axiovert 200M inverted fluorescence microscope with a 100x objective (N.A. 1.3). Images were analyzed using AxioVision software version 4.8.2.

### Immunofluorescence microscopy

All steps were performed at room temperature. Cells were harvested (800*g*, 10 min), washed once with PBS, fixed with 2% paraformaldehyde in PBS for 30 min and allowed to settle on slides for 30 min. Adhered cells were washed once with wash solution (0.1% normal goat serum in PBS) and permeabilized with 0.5% Trition X-100 for 30 min. Following permeabilization, cells were washed twice with wash solution and blocked with 10% normal goat serum (NGS) in PBS with 0.1% Triton X-100. Primary antibodies were diluted in block solution (C-myc, Thermo Fisher 9E10 1:500; *Tb*Aldolase 1:500; *Tb*HK 1:500; *Tb*FBP 1:1,000; *Tb*PFK 1:500) and incubated with cells for 1 h. Following primary antibody, slides were washed 3x with wash solution and incubated 1 h with secondary antibody (goat anti-mouse Alexa fluor 488 1:1,000, goat anti-rabbit Alexa fluor 568 or Alexa fluor 647 1:1,000); (Thermo fisher) in block. Wide-field images were taken using a Zeiss Axiovert 200M, 100x objective (N.A. 1.3), and analyzed using AxioVision software version 4.8.2. Super-resolution images were obtained using a Leica SP8X microscope equipped with HyD detector and 63x objective (N.A. 1.4).

### Electron microscopy processing and imaging

Cells were harvested (5×10^7^, 800*g* 10 min), washed three times with PBS, and fixed (2% paraformaldehyde, 2.5% glutaraldehyde in 100mM phosphate buffer pH 7.4). Cells were stored at 4°C for no longer than 2 days before being processed as described previously (*23*). Glycosome area measurements were performed using FIJI. Area of visible glycosomes for 15 fields was measured using the measure tool. To calculate the glycosome area as a percentage of cell area, the area of glycosomes from each cell was summed and divided by the total cell area visible.

### Biochemical analysis of glycosome protein localization

Cells were seeded at 1×10^5^ cells/ml and RNAi against *Tb*Pex13.2 was induced with 1 μg/ml doxycycline. After 4 days of induction, cells were harvested (800*g*, 10 min) and washed once with PBS. Cells (2×10^7^) were lysed using 1 volume wet weight silicon carbide abrasive and breakage confirmed by microscopy. Abrasive was removed by centrifugation (100*g*, 1 min) and supernatant was transferred to a new tube, followed by removal of nuclei (1,000*g*, 15 min). Supernatant was transferred to a new tube and the organelle-rich fraction was separated from the cytosolic fraction by centrifugation (17,000*g*, 15 min). Cytosolic proteins were precipitated with 4 volumes of acetone, incubated on ice for 1 h, and centrifuged at 17,000*g* for 10 min, 4°C. Pelleted organelle-rich fraction and cytosolic proteins were separated by SDS-PAGE and proteins detected by western blotting.

## RESULTS

### *Tb*Pex13.2 is expressed in BSF and PCF parasites

Antibodies raised against recombinant *Tb*Pex13.2 detected a protein of the expected size (~ 28 KDa) in PCF and BSF parasites, which was reduced upon induction of RNA interference (Fig. 1). These antibodies sometimes detected a larger ~50 kDa protein. This larger band was not reduced in *Tb*Pex13.2 RNAi cell lines, suggesting it was a non-specific, cross-reactive protein. To confirm that this upper band was not a dimer of *Tb*Pex13.2, we treated lysates with DTT (10 mM) and NEM (10 mM) to block free reactive thiols and prevent dimerization post-lysis. Neither treatment affected the abundance of the 50 kDa species (Fig. S1).

**Figure 1.**
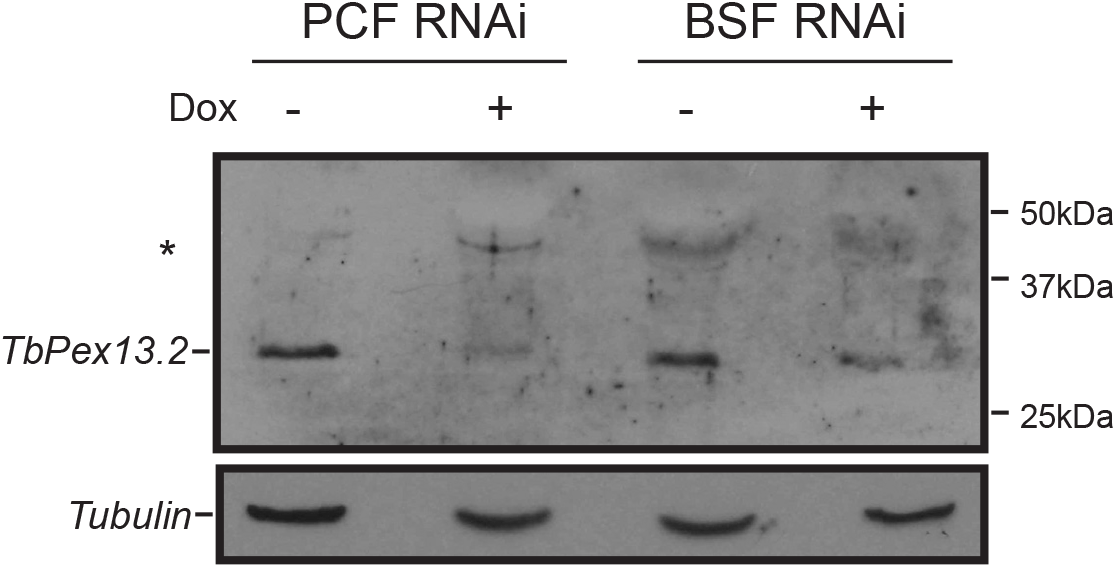
Antibodies raised against recombinant *Tb*Pex13.2 recognize a 28 kDa protein in PCF and BSF parasites that is reduced upon induction of RNAi. PCF and BSF parasites harboring pZJM:*Tb*Pex13.2 were grown for 4 days with and without doxycycline to induce RNAi. Lysates from 2×10^6^ cells were probed with antisera generated against r*Tb*Pex13.2 aa2-150. Tubulin was used as a loading control. *Larger cross-reactive band.

### *Tb*Pex13.2 is an integral glycosomal membrane protein with its N-terminus exposed to the cytosol

To confirm that *Tb*Pex13.2 was localized to glycosomes, we resolved organelles via density gradients and analyzed fractions by western blotting with antibodies against *Tb*BiP and *Tb*aldolase. Under our conditions, the ER is less dense than glycosomes and equilibrates at the top of the gradient. Consistent with being a glycosome protein, *Tb*Pex13.2 was detected in fractions 14-20 that also contained the glycosome protein aldolase (Fig. 2). In contrast, the ER protein, *Tb*BiP, was detected in fractions 20-32.

**Figure 2.**
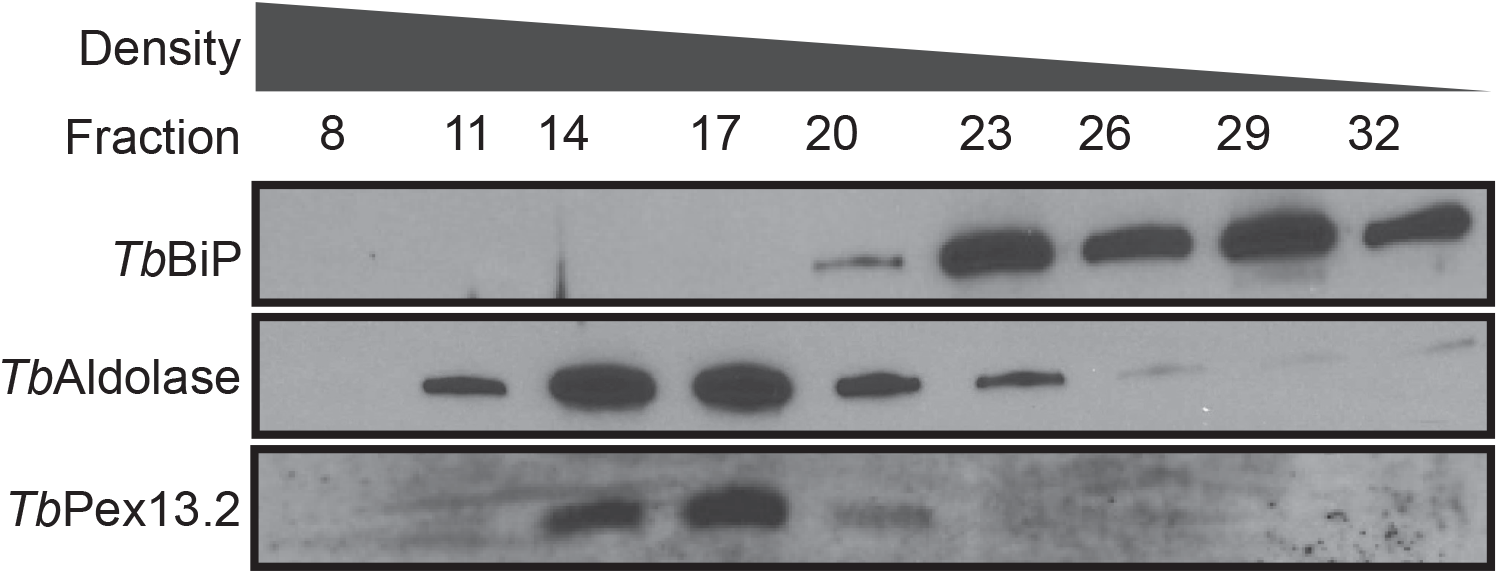
*Tb*Pex13.2 equilibrates with glycosome proteins in density gradients. Post-nuclear cell lysate was centrifuged through an Optiprep/sucrose gradient and 1ml fractions were collected from the bottom of the gradient. Protein was precipitated, separated by SDS-PAGE, and analyzed by western blot with antibodies against *Tb*aldolase, *Tb*BiP, and *Tb*Pex13.2.

*Tb*Pex13.2 is predicted to have at least 2 transmembrane (TM) domains (*14*, *24*). To confirm that *Tb*Pex13.2 is an integral membrane protein, we sequentially extracted membrane-enriched fractions with low-salt, high-salt, and sodium carbonate (Fig. 3A). The soluble glycosome matrix protein *Tb*aldolase was detected in the low and high-salt supernatants, whereas *Tb*Pex13.2 was detected only in the pellet following sodium carbonate extraction, as expected for an integral membrane protein. Orientation of the protein in the membrane was determined by protease protection assays with cells expressing *Tb*Pex13.2 fused to either an N-terminal myc epitope (myc*Tb*Pex13.2) or an N-terminal mycBirA* tag (mycBirA\**Tb*Pex13.2). Treatment of cells with proteinase K (PK) in the absence of Triton X-100 resulted in loss of myc signal, indicating that the N-terminus of the protein was accessible by PK and exposed to the cytosol (Fig. 3B). *Tb*aldolase is a matrix protein with a protease resistant core. Full-length *Tb*aldolase is detected in PK treatment without detergent indicating that glycosome integrity is not compromised during treatment. After treatment with PK and detergent, we observed a smaller proteolytic product as seen in previous studies documenting the protease-resistant nature of *Tb*aldolase (*25*). These results suggest that the N-terminus of *Tb*Pex13.2 is on the cytosolic side of the glycosome membrane.

**Figure 3.**
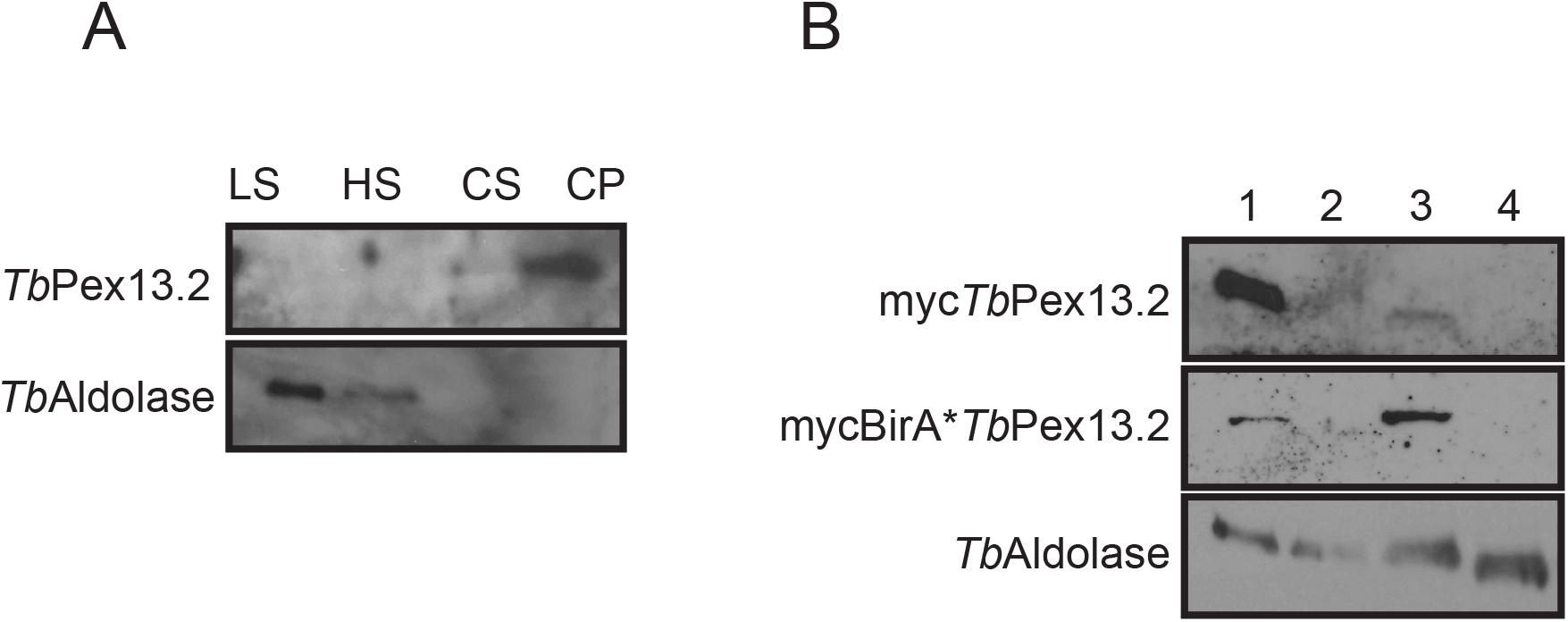
*Tb*Pex13.2 is an integral membrane protein with the N-terminus exposed to the cytosol. (A) Membrane protein extraction of whole cell membranes. Cells (10^7^) were sequentially extracted with low-salt, high-salt, and sodium carbonate. Proteins were precipitated from supernatants (LS, HS, CS) and final carbonate pellet (CP), separated by SDS-PAGE, transferred to nitrocellulose, and analyzed by western blotting with antibodies against *Tb*Pex13.2 and aldolase. (B) Protease protection assay. Cell lysates (lane 1) were treated with protease K (lane 2), Triton X-100 (lane 3), or Triton X-100 and protease K (lane 4). Samples were resolved by SDS-PAGE and analyzed by western blots with anti-myc antibodies.

### *Tb*Pex13.2 is localized to discrete foci in glycosomes

Antibodies raised against recombinant *Tb*Pex13.2 do not work in immunofluorescence assays (IFA). Because of this, we analyzed cell lines expressing *Tb*Pex13.2 fused to an N-terminal myc tag (myc*Tb*Pex13.2) (Fig. 4 A, B). Anti-myc antibodies labeled punctate structures that were distributed throughout the cell and exhibited limited overlap with the glycosome marker *Tb*aldolase. We used Mander’s overlap coefficients (MOCs) to quantify the extent to which myc*Tb*Pex13.2 co-localized with the glycosome marker aldolase and the ER marker *Tb*BiP (Fig. 4 C, D). MOC values range from 0 (no overlap) to 1 (complete overlap) (*26*). MOC for *Tb*aldolase and myc*Tb*Pex13.2 was 0.62 ± 0.16 suggesting limited co-localization. To better define the relationship between myc*Tb*Pex13.2 and *Tb*aldolase staining, we calculated Mander’s M1 and M2 values, which reveal the extent to which pixels in one channel overlap with the other. The M1 and M2 values were 0.49 ± 0.20 and 0.91 ± 0.10, respectively. These numbers indicate that 49% of the signal in channel 1 (*Tb*aldolase) overlaps with channel 2 (myc) and that 91% of the signal in channel 2 (myc) overlaps with channel 1 (*Tb*aldolase). These results suggest that myc*Tb*Pex13.2 is limited to a restricted portion of the glycosome. MOC values for *Tb*BiP and myc*Tb*Pex13.2 was 0.17 ± 0.07, suggesting that these two proteins do not co-localize. In agreement with the MOC, M1 and M2 values, we observed that myc*Tb*Pex13.2 was not evenly distributed throughout glycosomes but was restricted to a few distinct foci within each organelle. Thirty-seven percent of the myc*Tb*Pex13.2-positive glycosomes had one foci, 35% had two foci, and 24% had 3 or more. Of *Tb*aldolase positive glycosomes, approximately 4% had no visible myc*Tb*Pex13.2 foci (Fig. 4E).

**Figure 4.**
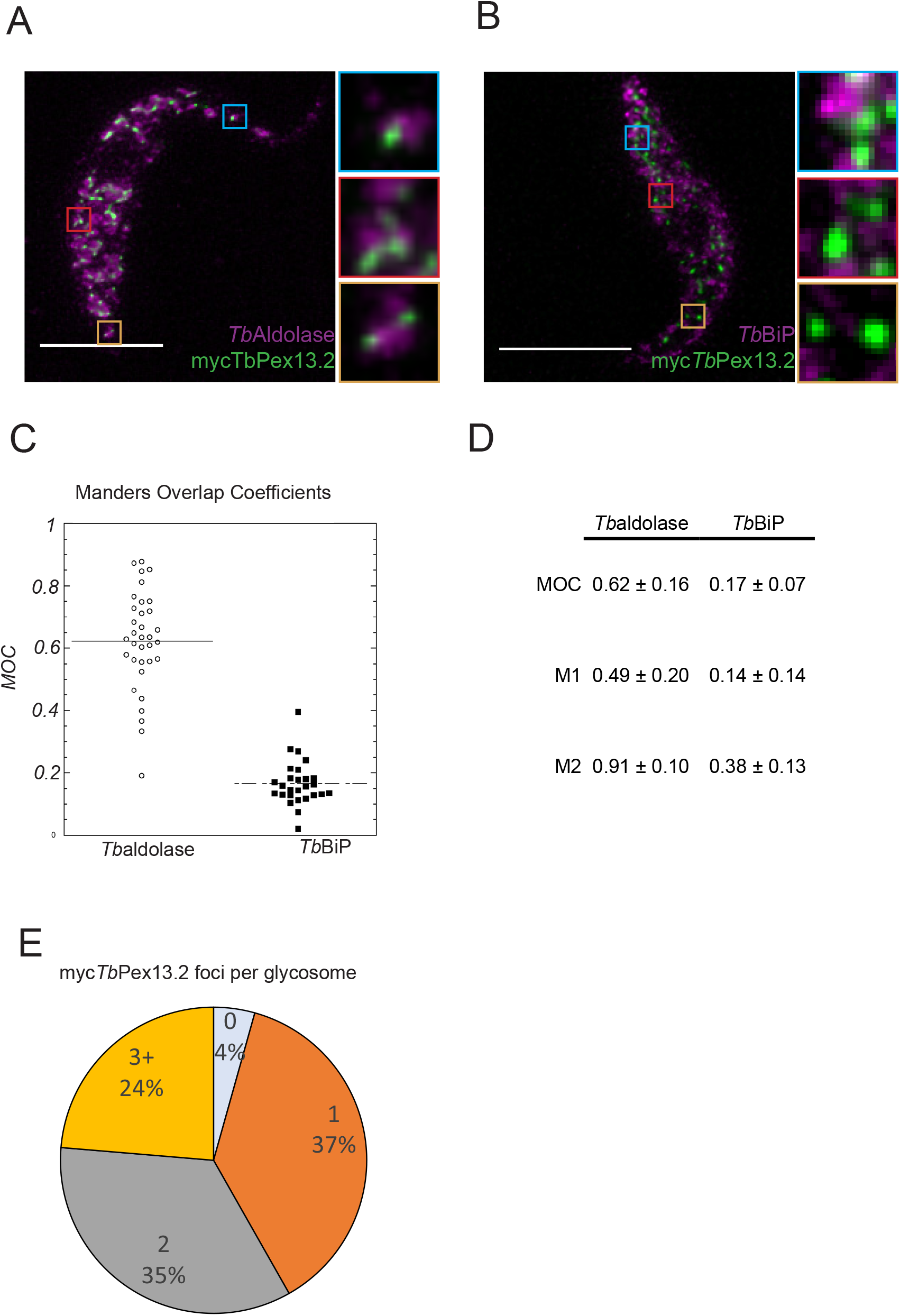
Epitope-tagged myc*Tb*Pex13.2 localizes to glycosomes. (A,B) Cells were fixed, permeabilized, and stained with antibodies against myc (green) and *Tb*aldolase or *Tb*BiP (magenta). Representative images of maximum intensity projections of deconvolved Z-stacks are shown. Scale bar indicates 5 μm. (C) Plots of Mander’s overlap coefficients (MOC) and Manders M1 and M2 values were calculated from individual cells using Z-stacks with the Mander’s coefficient plugin in FIJI. Each point represents analysis of one cell. Myc*Tb*Pex13.2 and *Tb*Aldolase n=34. myc*Tb*Pex13.2 and *Tb*BiP n=27. (D) MOC, M1 and M2 values. (E) Pie chart representing the counted number of myc*Tb*Pex13.2 foci per organelle.

### Silencing of *Tb*Pex13.2 in PCF parasites reduced import efficiency of PTS2 proteins

Prior work revealed that silencing *Tb*Pex13.2 in BSF parasites was detrimental (*14*). At that time PCF RNAi cell lines could not be established. We were able to silence *Tb*Pex13.2 in both PCF and BSF parasites via inducible RNA interference and western analysis indicated that *Tb*Pex13.2 levels were reduced ~95% upon induction of RNAi (Fig. 1). Silencing of *Tb*Pex13.2 did not alter the overall expression levels of the glycosome matrix protein *Tb*aldolase, or the glycosome membrane proteins *Tb*Pex13.1 or *Tb*Pex14 (Fig. 5A) in BSF or PCF parasites. While we observed no change in growth rate after induction of RNAi in PCF parasites (Fig. 5B), BSF growth was slowed upon induction of silencing (Fig. 5C) as reported previously (*14*).

**Figure 5.**
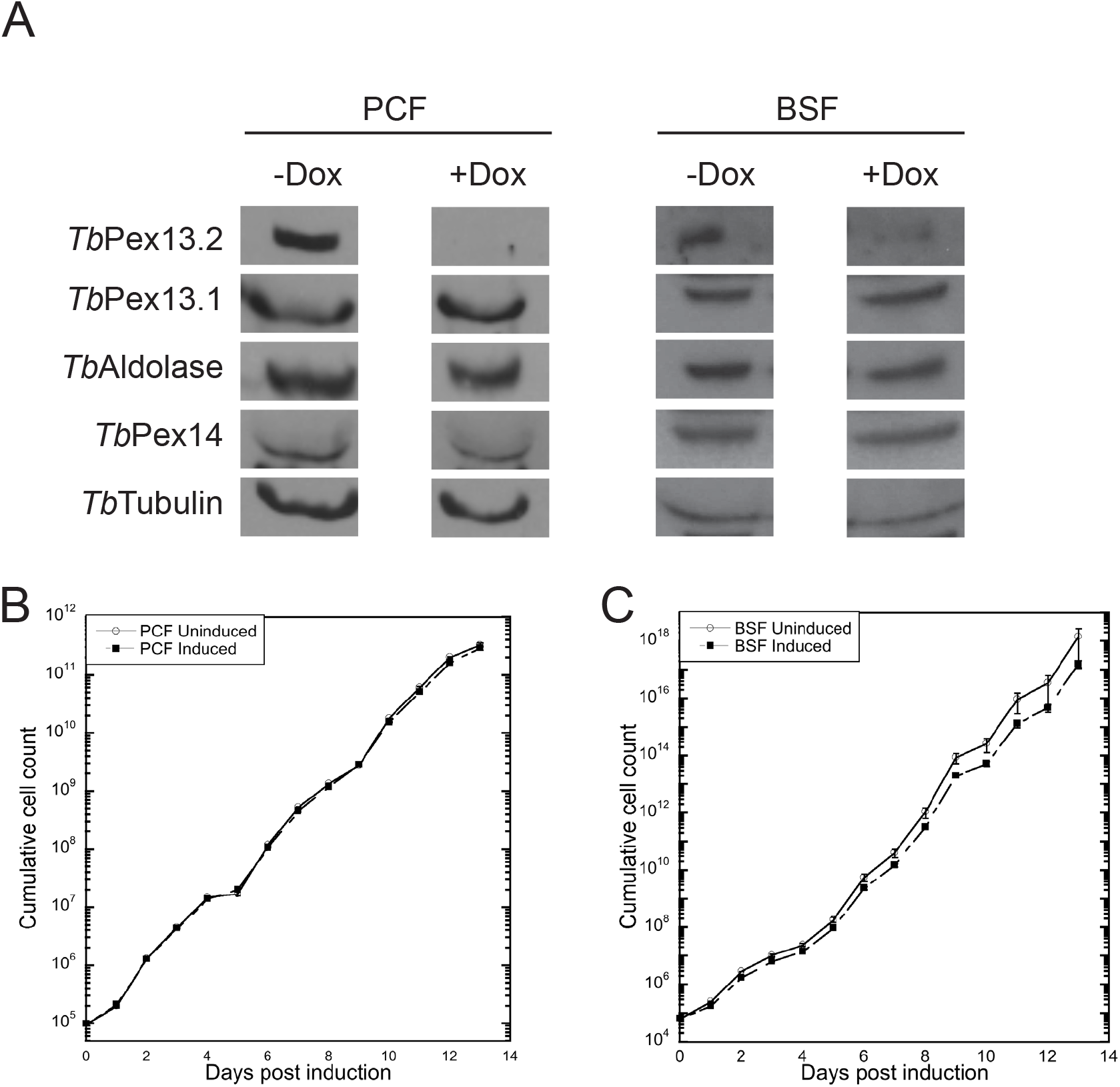
Silencing of *Tb*Pex13.2 does not influence expression of other glycosome proteins and slows BSF growth. (A) Cultures were grown for 96 h and 5×10^6^ cells were analyzed by western blotting with antibodies against *Tb*Pex13.2, *Tb*Pex13.1, *Tb*aldolase, and *Tb*Pex14. Tubulin was used as a loading control. Growth curves for PCF (B) and BSF (B) parasites grown with and without doxycycline.

In higher eukaryotes, Pex13 functions in peroxisomal protein import. We used immunofluorescence assays (IFA) to determine if *Tb*Pex13.2 silencing altered glycosome protein import in PF parasites. In uninduced cells, the glycosome proteins *Tb*aldolase, *Tb*HK, *Tb*FBP and *Tb*PFK all localized to punctate structures characteristic of glycosomes (Fig. 6A). After 4 days of induction, IFA revealed an increase in the cytosolic localization of *Tb*aldolase. In contrast, the localization of *Tb*HK, *Tb*FBP, and *Tb*PFK was not altered in these cells. Under standard culturing conditions in SDM79 media (5 mM glucose), the mislocalization of glycosome proteins is generally lethal. Previous work has shown that removal of glucose from the media rescues this phenotype (*27*). We measured glycosome protein localization in low-glucose media (SDM79θ, 5 μM glucose) with the expectation that under these conditions we could score phenotypes that would be lethal in cells grown in SDM79. When cells were grown in SDM79θ, tet-induced mislocalization of *Tb*aldolase was increased. In contrast, even in low-glucose conditions, the *Tb*HK, *Tb*FBP and *Tb*PFK localization was not altered upon *Tb*Pex13.2 silencing.

**Figure 6.**
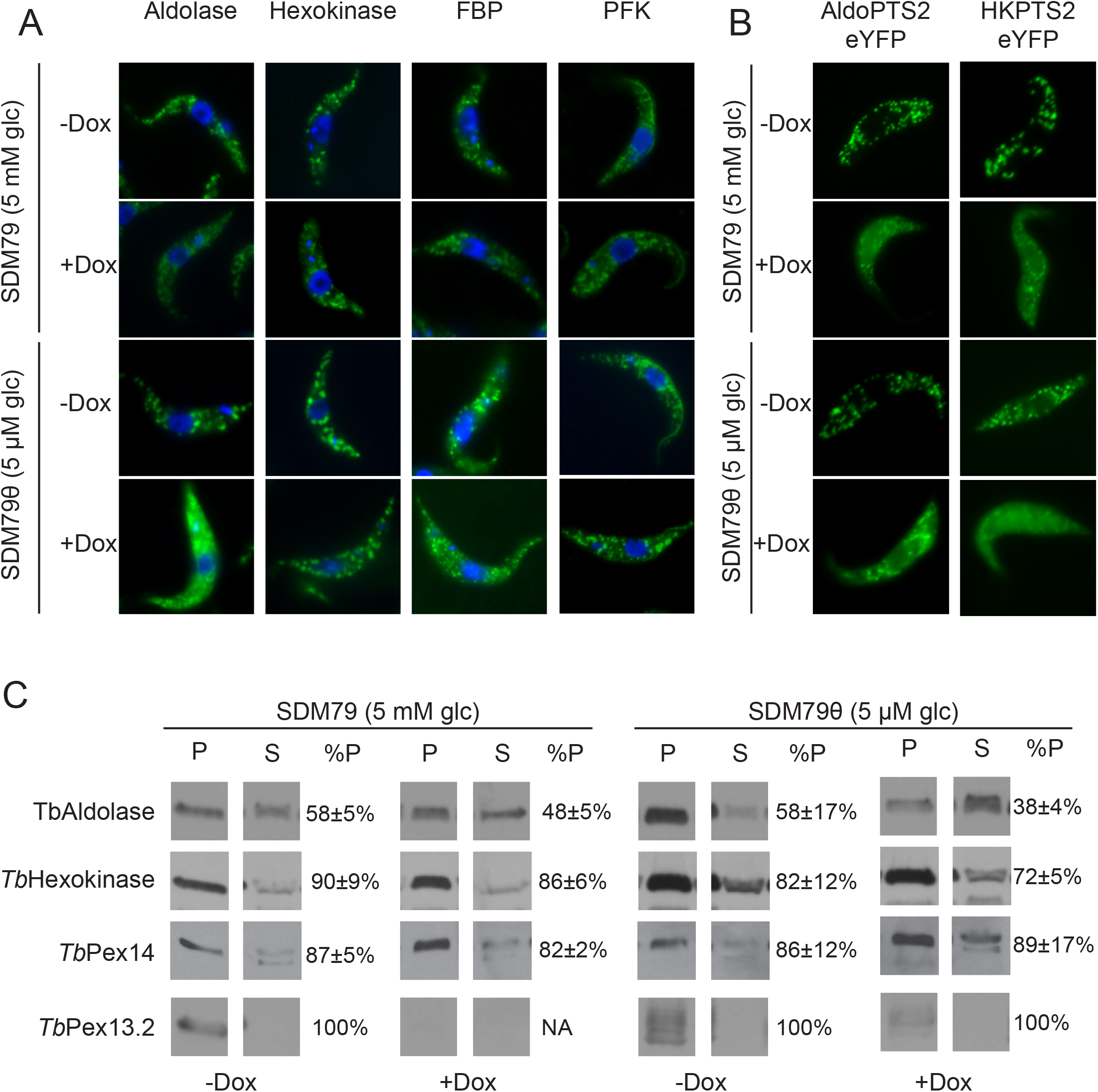
Silencing of *Tb*Pex13.2 results in mislocalization of a subset of glycosome proteins. (A) IFA assays of PCF cells grown in the presence or absence of doxycycline (DOX). Cells were labeled with antibodies against *Tb*aldolase, *Tb*hexokinase, and *Tb*fructose 1,6-bisphosphatase (*Tb*FBP) and detected using goat anti-rabbit secondary antibodies conjugated with Alexa Fluor 488 (Green). Nuclear and kinetoplast DNA were stained with DAPI (Blue). (B) Live cell microscopy of cells constitutively expressing eYFP fused with the peroxisome targeting sequence 2 (PTS2) of aldolase or hexokinase (Green). (C) Biochemical analysis of mislocalization. Cells were lysed by silicon carbide abrasive and were centrifuged to obtain an organelle enriched pellet. Cytosol proteins in supernatant were precipitated using 4 volumes of acetone. Samples were separated by SDS-PAGE and analyzed by western blot. Mislocalization was quantified by densitometry of three biological replicates using ImageJ.

To quantify import efficiencies of each protein, we used western analysis to determine the percent of protein associated with organelle fractions in uninduced and induced *Tb*Pex13.2 RNAi cell lines. Cells were lysed and centrifuged to obtain a membrane-rich fraction and western analysis used to calculate the percentage of each protein associated with membrane (glycosomes) and soluble (cytosolic) fractions (Fig. 6C). In uninduced cells, 58 ± 5% of *Tb*aldolase was associated with the pellet compared to 48.5 ± 5% in induced cells. This mislocalization was increased in low-glucose media with 58.5 ± 5% associated with the pellet in uninduced cells and 38 ± 4% in *Tb*Pex13.2 silenced cells. In contrast to *Tb*aldolase, *Tb*FBP, *Tb*PFK, and *Tb*HK localization did not change significantly when *Tb*Pex13.2 was silenced in high or low-glucose media. While both *Tb*HK and *Tb*aldolase have type 2 peroxisomal targeting sequences (PTS2s), only *Tb*aldolase localization was affected by *Tb*Pex13.2 silencing. To assess the extent to which each PTS2 sequence contributed to this *Tb*Pex13.2-dependent localization, we followed the localization of eYFP fused to either the PTS2 of *Tb*aldolase or *Tb*HK. Fluorescence microscopy of living cells revealed that both *Tb*AldoPTS2eYFP and *Tb*HKPTS2eYFP were mislocalized to the cytoplasm in *Tb*Pex13.2 deficient cells (Fig. 6B).

We next used transmission electron microscopy (TEM) to determine how depletion of *Tb*Pex13.2 altered overall glycosome morphology (Fig. 7A). Under these conditions, glycosomes are spherical, electron dense organelles. In SDM79 media, glycosomes comprised 3.67% ± 1.76% and 2.14% ± 0.84% of the cell volume in wild-type or *Tb*Pex13.2-deficient cells, respectively. In SDM79θ media, glycosomes comprised 6.18% ± 1.75% and 5.77% ± 2.11% of the cell volume in wild-type or *Tb*Pex13.2-deficient cells, respectively. These results suggest that glycosomes morphology was not dramatically altered upon reduction of *Tb*Pex13.2 levels.

**Figure 7.**
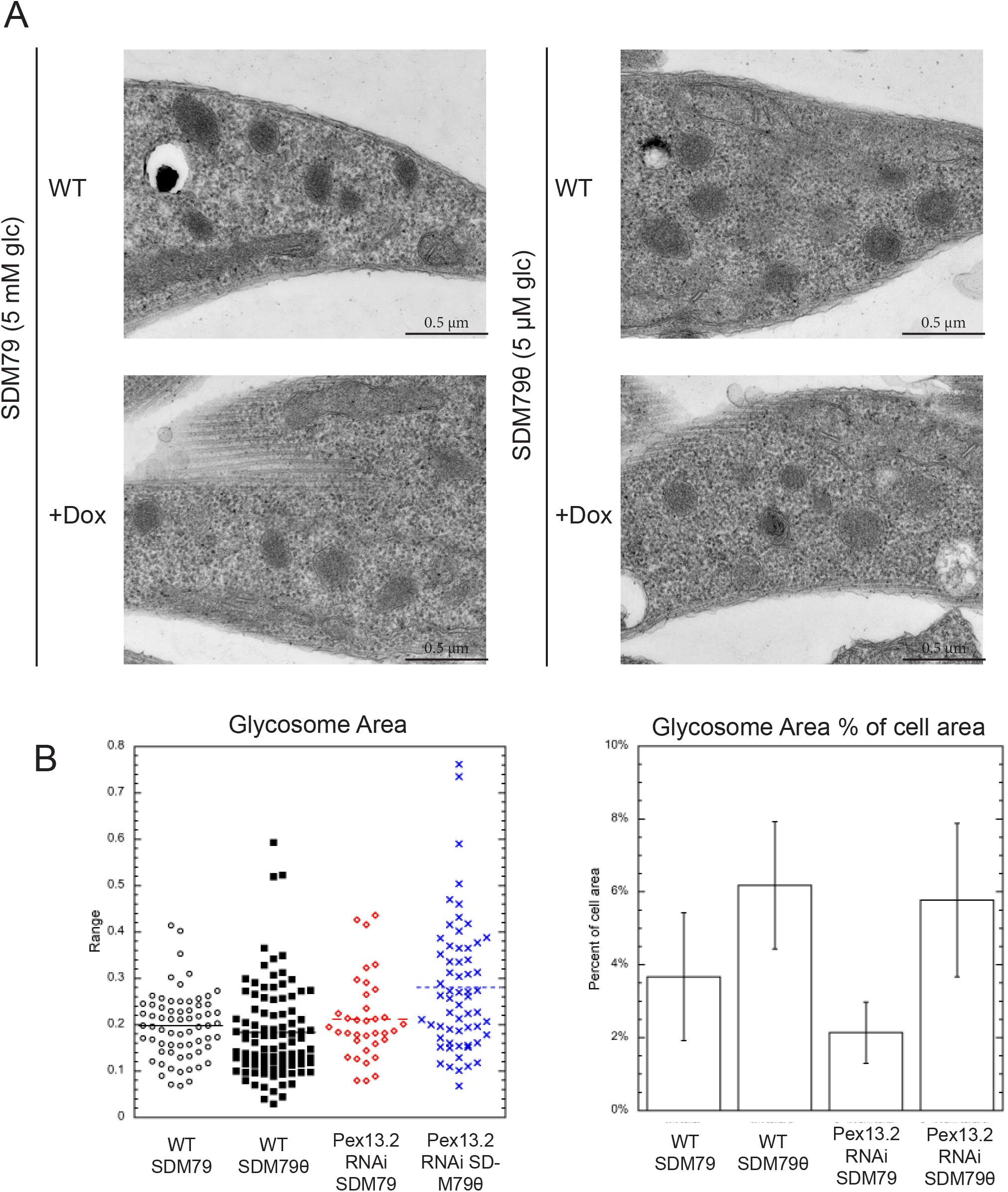
Electron microscopy of *Tb*Pex13.2 deficient cells. (A) EM analysis of wildtype and cells induced for RNAi (B) Analysis of glycosome size and area as a percentage of cell area. Twenty-five fields and at least 36 glycosomes per condition were measured.

**Figure 8.**
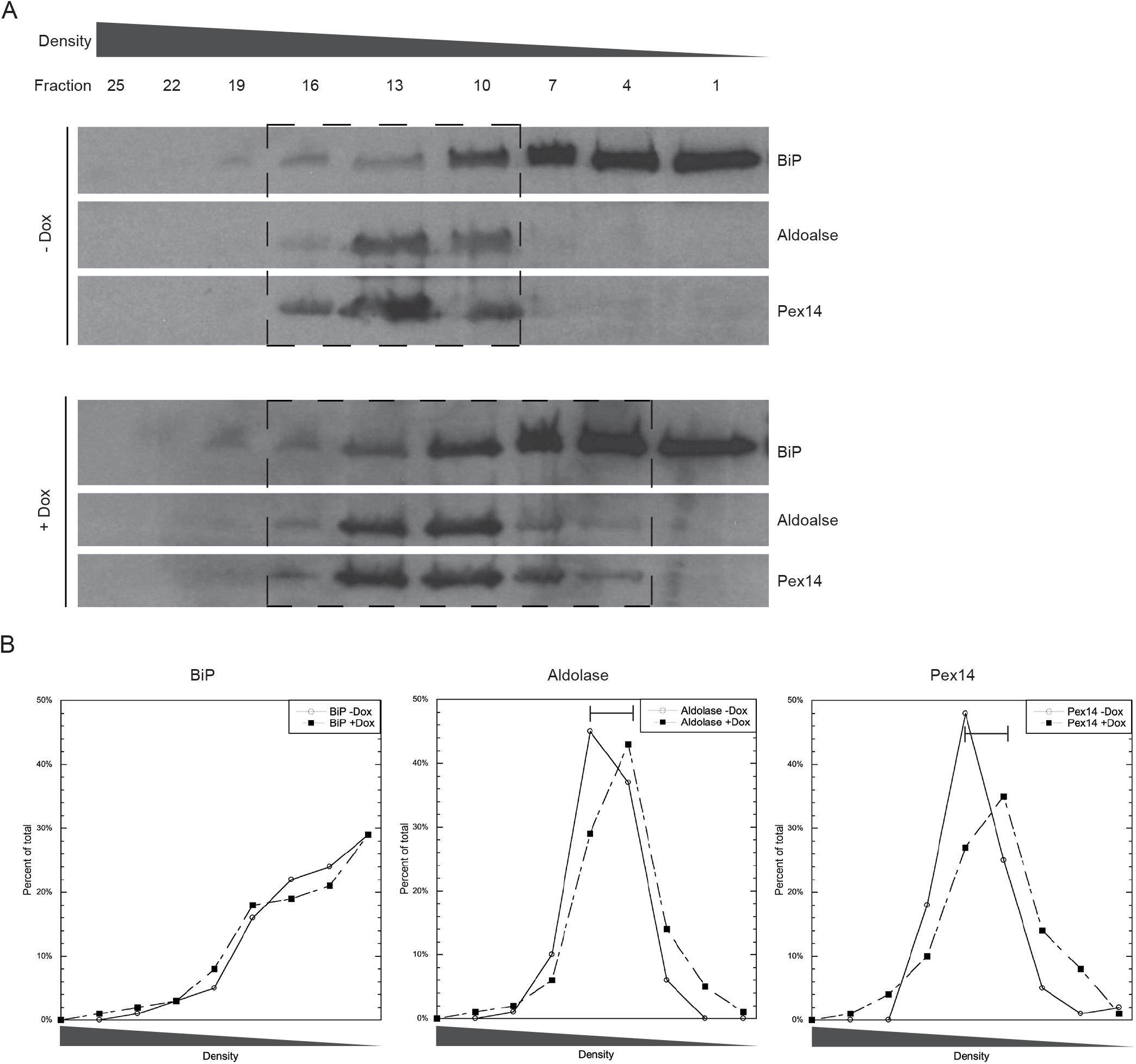
Glycosomes from *Tb*Pex13.2-deficient cells are lighter than wild-type. (A) Cells (5×10^8^) were harvested, lysed mechanically with silicon carbide, and the glycosome-enriched pellet resolved on a linear Optiprep gradient. Fractions were taken from the top of the gradient and analyzed by western blotting. Boxes indicate glycosome fractions (B) Western blot bands were quantified using densitometry and the percentage of protein in each fraction was plotted. Presented is a representative gel from three biological replicates.

We reasoned that *Tb*Pex13.2 silencing may interfere with the import of proteins for which we have no antibodies. Additionally, depletion of a protein localized to the glycosome membrane may affect other biochemical properties such as density. To assess this, we resolved organelles on a sucrose/Optiprep gradient and followed sedimentation via western analysis of glycosomal proteins. In uninduced cells, *Tb*aldolase and *Tb*Pex14 were detected in fractions 10-16 as is typical for glycosome proteins. In induced cells, however, these proteins were detected in higher fractions (4-16) suggesting that glycosomes in *Tb*Pex13.2 deficient cells are lighter than those isolated from parental cells. In contrast, western analysis with anti-*Tb*BiP antibodies revealed that the sedimentation of the ER was not altered in *Tb*Pex13.2 deficient cells.

## DISCUSSION

Kinetoplastids harbor specialized peroxisomes called glycosomes that are essential to parasite survival. The majority of the glycolytic pathway is compartmentalized within these organelles and disruption of compartmentalization is lethal under standard conditions (*5*, *8*, *27*–*29*). As glycosomes are essential organelles in kinetoplastids, the mechanisms for their maintenance are attractive drug targets (*4*, *30*, *31*).

Like peroxisomes, glycosomes do not have DNA and all matrix proteins are imported post-translationally in a process that is fairly well-conserved. Glycosome and peroxisome matrix proteins are synthesized in the cytoplasm where they are identified by the soluble receptors, Pex5 and Pex7, which recognize peroxisomal targeting sequence (PTS) 1 and PTS2, respectively (*8*). The receptor:cargo complex is then targeted to the receptor docking complex on the glycosome/peroxisome membrane containing the proteins Pex13 and Pex14 (*7*, *12*, *32*). *T. brucei* is unusual in that it has two Pex13s, which have been named *Tb*Pex13.1 and *Tb*Pex13.2 (*14*, *15*). The two proteins share low overall sequence identity with each other or with other Pex13s and are conserved in *T. brucei, T. cruzi* and *Leishmania*. *Tb*Pex13.1 was the first to be identified and characterized in *T. brucei*. Fusions with green fluorescent protein localized to glycosomes and silencing *Tb*Pex13.1 resulted in glycosome protein mislocalization and cell death. *Tb*Pex13.2 was identified later and silencing of this protein in BSF parasites resulted in glycosome protein mislocalization and cell death. These findings indicate that the two proteins are not redundant, however, the specific role that each plays in glycosome biogenesis is unclear.

Here, we report the characterization of *Tb*Pex13.2 and a phenotypic characterization of RNA interference cell lines in which *Tb*Pex13.2 is silenced. In agreement with previous work, we found that silencing *Tb*Pex13.2 slowed the growth rate of BSF parasites (*14*). While the effect we observed was not as dramatic as previous work, this is likely due to variation in penetrance. As previously observed with the tagged *Tb*Pex13.2 (*14*), we found that native *Tb*Pex13.2 was an integral glycosome membrane protein. Furthermore, we were able to demonstrate through protease protection assays that the N-terminus of *Tb*Pex13.2 is exposed to the cytosol where it can interact with proteins involved in protein import. Unfortunately, we could not express C-terminally tagged *Tb*Pex13.2 in *T. brucei* and were not able to determine if the C-terminus is localized to the cytosolic or matrix side of the glycosome.

We found that *Tb*aldolase, which has a PTS2 sequence was mislocalized in *Tb*Pex13.2-deficient parasites while the import of PFK harboring a PTS1 was not affected. During the import process, Pex7 binds to the N-terminus of Pex13 that is comprised of a YG rich sequence in S. *cerevisiae* and *A. thaliana* (*33*, *34*). Both *Tb*Pex13.1 and *Tb*Pex13.2 have a similar YG rich domain in their N-termini. It may be that depletion of *Tb*Pex13.2 reduces the number of Pex7 binding sites at the glycosome membrane resulting in reduced PTS2 import while having minimal effect on protein import of PTS1 proteins such as PFK. *Tb*FBP import was not affected in *Tb*Pex13.2-deficient cells, however, this protein has both PTS1 and PTS2 sequences and may be imported by a combination of PTS1 *and* PTS2 import pathways.

In previous studies using BSF parasites, digitonin extractions revealed that silencing *Tb*Pex13.2 impaired import of both PTS1 and PTS2 proteins. So far, we have not observed defects in PCF PTS1 import. This may be because we followed the import of a different set of proteins due to antibody availability. Additionally, it may be that the role of *Tb*Pex13.2 in protein import varies with lifecycle stage. Finally, the observed differences may be a consequence of the methods used to assess import. Previous studies utilized digitonin fractionations while we measured import via IFA and quantification of organelle associated proteins.

The selective defect in aldolase (PTS2) protein import may be a reflection of the different mechanisms that direct PTS2 and PTS1 protein import. The receptor Pex7, unlike Pex5, is necessary but not sufficient for PTS2 protein import and several PTS2 co-receptors have been described. These co-receptors are species-specific and include Pex18 and Pex21p in *Saccharomyces cerevisiae* (*35*), Pex20p in *Yarrowia lipolytica* (*36*), and the long isoform of Pex5 (Pex5L) in mammals (*37*). All three co-receptors have Pex7-binding domains. *T. brucei* has two isoforms of Pex5, however, the functional significance of the two forms are very different from what is observed in mammals. Whereas, *Hs*Pex5L is a splicing variant with an internal Pex7BD, *Tb*Pex5L is the full-length protein while the shorter form is a result of a proteolytic cleavage (*38*). Therefore, a co-receptor has not yet been identified in *T. brucei*. It is attractive to speculate that *Tb*Pex13.2 may be playing this role in kinetoplastids. Indeed, the N-terminus containing the YG-rich region faces the cytoplasm and is physically positioned to function in this capacity.

We were surprised to find that *Tb*aldolase and *Tb*HK, which both have PTS2s, behaved differently in *Tb*Pex13.2 deficient cells. *Tb*aldolase import was disrupted while *Tb*HK was not. We imagine two possible scenarios for these results. It is possible that not all PTS2s are with the same efficiency. Alternatively, there may be information outside of the PTS2 sequence that influences the glycosome localization of *Tb*HK. To discern which scenario was at play, we fused the PTS2 of *Tb*HK and the PTS2 of *Tb*aldolase to enhanced yellow fluorescent protein (eYFP) and found that the import of each is disrupted in *Tb*Pex13.2-deficient cells. This finding indicates *Tb*Pex13.2 is required for efficient import of both PTSs and reveals that that *Tb*HK contains information for glycosomal localization outside of the PTS2 sequence.

The formation of myc*Tb*Pex13.2 foci on the glycosome membrane was interesting because of its similarity to the localization of other import process-related peroxins in other systems. Recent super-resolution studies of peroxisomes have provided insight into the heterogeneous distribution of different peroxins within peroxisomes (*39*). Membrane-bound Pex5 and Pex14 generally colocalized in structures that range in shape from round to elliptical, ring-like structures. Pex11, which functions in peroxisome proliferation and not protein import, and a soluble matrix protein SCP2 also exhibit heterogenous distribution. The functional relevance of these distribution patterns is unclear; however, the use of super-resolution imaging is essential to resolving the mechanism and consequence of such localization patterns. To our knowledge, this is the only example of heterogenous intra-glycosomal localization of peroxins in parasites and future studies are necessary to understand the significance of such localizations.

## Acknowledgements

This work was funded by an NSF COBRE P20 grant P20GM109094. We thank Clemson Light Imaging Facility for imaging assistance and use of equipment and Dr. Jay Bangs (University of Buffalo, Buffalo, NY), Dr. Christine Clayton (Zentrum für Molekulare Biologie der Universität Heidelberg, Germany), and Dr. Paul Michels (University of Edinburgh, UK) for antibodies. We also thank Wandy Beatty (Washington University in St. Louis) for contributing to the transmission electron microscopy work.

## Figure Legends

**Figure S1.**
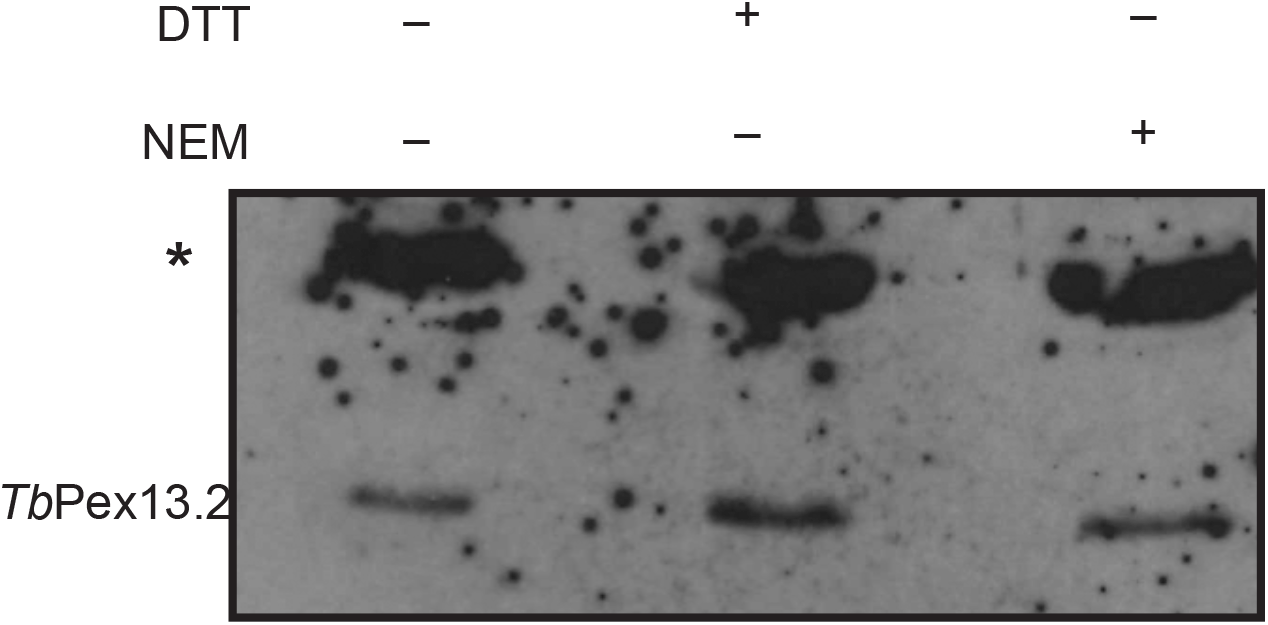
Antibodies raised against recombinant *Tb*Pex13.2 recognize a 28 kDa protein in PCF and BSF parasites that is reduced upon induction of RNAi. Higher molecular weight bands are not multimers of *Tb*Pex13.2. Bloodstream form (10^6^) lysates were treated with DTT to reduce disulfide bonds and NEM to block free reactive thiols to prevent dimerization post lysis. Lysates were resolved by SDS-PAGE and western blots probed with α-*Tb*Pex13.2 antibodies. Asterisk indicates non-specific cross-reactive band.

